# Catalytically inactive dKbCas12d guided by sgRNA and new insights into its binding through a single-molecule approach

**DOI:** 10.1101/2023.12.12.571249

**Authors:** Aleksandr Alekseev, Valerii Klimko, Marina Abramova, Irina Franzusova, Victor Vinnik, Aleksandra Vasileva, Polina Selkova, Mikhail Khodorkovskii, Anatolii Arseniev

**Affiliations:** Peter the Great St. Petersburg Polytechnic University, 195251 Saint Petersburg, Russia; Complex of NBICS Technologies, National Research Center “Kurchatov Institute”, 123182 Moscow, Russia

## Abstract

CRISPR-Cas12d is a distinct V-D type system discovered in the metagenomes of Candidate Phyla Radiation bacteria. It stands out from most closely related systems due to its 17-19 nucleotide short spacer region and specialized stabilizing scoutRNAs. We made significant improvements to this system by modifying its scoutRNA to create sgRNA, which greatly simplifies its use. We found mutations in the RuvC domain of the effector protein KbCas12d that resulted in loss of nuclease activity. We obtained two catalytically inactive dKbCas12d variants: D827A and E913A. Using the optical tweezers technique, we demonstrated the high specificity of dKbCas12d in binding targets on individual DNA molecules. Engineered sgRNA and catalytically inactive dKbCas12d variants have promising applications in biotechnology for the precise regulation of gene expression and molecular diagnostics.

## INTRODUCTION

CRISPR-Cas systems are defense mechanisms used by bacteria and archaea to provide adaptive immunity against various mobile genetic elements such as transposons, bacteriophages, and plasmids [1,2]. The operation of these systems is based on the action of RNA-protein complexes consisting of Cas effectors and guide RNAs. Complementary pairing of guide RNAs with the foreign genome enables Cas effectors to precisely recognize and cleave specific sites in the invader’s nucleic acid, halting further infection. For genetic engineering purposes, Cas effectors can be directed towards desired sites in the genomes of prokaryotic and eukaryotic organisms by utilizing guide RNAs to selected sequences [3,4]. Characterization of novel CRISPR-Cas proteins allows us to expand the range of potential genome engineering tools and provides fundamental insights into the operation of bacterial defense systems [5,6].

CRISPR-Cas12d (previously named CRISPR-CasY) systems were discovered by studying metagenomic data from the Candidate Phyla Radiation (CPR) bacteria [6,7]. CPR bacteria are not culturable in the laboratory and represent more than 15% of the total bacterial diversity. [7–9]Presumably symbionts of other prokaryotes, these organisms are characterized by a compact cell size and small genomes. CRISPR-Cas12d systems found in these bacteria are also characterized by a small size of effector genes [10].

The locus of the CRISPR-Cas12d system includes a CRISPR array, the *cas1* gene, which is thought to be responsible for adaptation, and the *cas12d* gene, which encodes a nuclease of about 1200 amino acids. [6,7]The sequences between the direct repeats of the CRISPR array are also reduced and represent short spacers of 17-19 nucleotides in length. For KbCas12d (also named CasY.1) system discovered in Katanobacteria has been previously shown that cells carrying the CRISPR-Cas12d locus are capable of interference defense against foreign mobile elements *in vivo* [7]. Later research discovered [11] that CRISPR-Cas12d systems require additional scoutRNA. *In vitro* experiments mainly focused on the CRISPR-CasY15 system, observed the presence of a 5’-TR-3’ – PAM sequence, necessity of scoutRNA for specific nuclease activity, collateral activity, and strong specificity of the system with spacer sequence substitutions. Similar to other type V systems, the RuvC domain which is responsible for DNA cleavage has been identified [5,10,12,13].

In this study, we aimed to identify mutations that would disable the active site of the Cas12d effector protein KbCas12d. To achieve this goal, we produced recombinant wild-type and mutant proteins and compared *in vitro* activity. Furthermore, in order to effectively utilize the system, we fused crRNA and scoutRNA molecules to obtain single guide RNA. Since standard biochemical methods do not always provide a comprehensive assessment of DNA-protein interactions, we employed optical tweezers to gain a better understanding of the interaction between the catalytically inactive mutant and individual DNA molecules. This approach, along with the use of fluorescence imaging, allows for more direct visualization of specific, targeted DNA-protein interactions.

## MATERIALS AND METHODS

### Recombinant protein expression and purification

The wild-type and mutant forms dCas12d D827A and dCas12d E913A genes were cloned into pET21a plasmids under the control of the T7 promoter. His and MBP tags were added to the C-termini of the proteins in the cloning state for subsequent affinity purification. The resulting plasmids pET21_Cas12d, pET21_dCas12d_D827A, and pET21_dCas12d_E913A were transformed into competent *E. coli* Rosetta cells. A 10 ml overnight culture of transformed bacteria was diluted into 500 ml LB medium with ampicillin and cultured at 37°C until reaching an optical density of 0.6 OD. Expression of effector proteins was induced by adding 1 mM IPTG, followed by incubation at 16°C for 16 hours. Cells were then lysed in 15 ml lysis buffer (50 mM Hepes-HCl pH = 7.5, 500 mM NaCl, 10% glycerol, and 1 mM beta-mercaptoethanol) with the addition of 15 mg lysozyme. After sonication and centrifugation, the supernatant was filtered (0.22 μm pore size) and applied to a HisTrap HP 1 mL column (GE Healthcare) for affinity chromatography. After washing with lysis buffer containing 20 mM imidazole, the proteins were eluted with lysis buffer containing 300 mM imidazole. Fractions were concentrated using Amicon Ultra-4 centrifugal units (Merck Millipore) with a 100 kDa cutoff to a volume of 600 μl. At this stage, TEV protease was added to the samples to cleave MBP tag, and incubated overnight at 10°C. The resulting sample was applied to a Superdex 200 Increase 10/300 GL gel filtration column (GE Healthcare) equilibrated with 50 mM Hepes-HCl (pH = 7.5), 500 mM NaCl, 1 mM DTT, and 10% glycerol. Fractions containing monomeric protein were collected, concentrated, frozen in small volumes in liquid nitrogen and stored at -80°C.

### RNA synthesis *in vitro*

RNA synthesis was performed according to the HiScribe SP6 RNA Synthesis Kit protocol (NEB). Double-stranded DNA sequences containing SP6 promoter 5′ATTTAGGTGTGACACTATAG 3′ were used as DNA matrixes for RNA transcription.

G is the first base of the RNA tanscript (in supplementary files RNA sequences are indicated from the second base after G).

Fluorescently labeled RNA was also synthesized according to the HiScribe SP6 RNA Synthesis Kit protocol for the production of labeled transcripts using FAM-UTP nucleotides (Lumiprobe) in a 3:4 ratio to unmodified UTPs.

### Detection of nuclease activity *in vitro*

DNA cleavage reactions were performed using recombinant KbCas12d protein and linear dsDNA target (818 b. p.) obtained by the PCR method from plasmid pUC19 containing the native spacer of the KbCas12d system. The reaction was mix using 1× CutSmart buffer (50 mM potassium acetate, 20 mM Tris-acetate, 10 mM magnesium acetate, and 100 μg/mL BSA; NEB) and 0.5 mM DTT (1,4-dithiothreitol) with varying concentrations of recombinant KbCas12d or its mutant versions and RNA (as indicated in the results). The samples were then incubated at 37°C for 30 minutes. To stop the reaction, 4× loading dye containing 10 mM Tris-HCl (pH = 7.8), 40% glycerol, 40 mM ethylenediaminetetraacetic acid, 0.01% bromophenol blue, and 0.01% xylocyanol was added. Electrophoretic analysis of the reaction products was performed on either 1.5% agarose gels or 5% denaturing gels (urea 8M).

### Detection of binding activity *in vitro*

Electrophoretic mobility shift assay (EMSA) was utilized to assess the DNA-binding activity of dCas12d D827A and dCas12d E913A. The reactions were performed as described previously [14] on a DNA matrix obtained by amplification of the GRIN2B gene with a length of 132 bp limited by a primer containing a Cy-3 tag. RNP complexes consisting of protein and RNA in a 1:5 ratio in different ratios (1:1; 1:3; 1:10) relative to 20 nM matrix DNA were incubated for 30 min at 37°C. RNP complexes containing non-target RNA were used as negative controls. Samples were then subjected to a 5% PAGE gel to visualize binding.

### Optical tweezers setup

For single-molecule manipulations a custom-built dual trap optical tweezers setup based on the Axioimager.Z1 microscope was used as described previously [15,16]. In brief, two independent optical traps were generated using a Nd:YVO4 1064 nm CW laser (5 W, Spectra Physics BL-106C) and an oil immersion lens with a high numerical aperture. Manipulations with single DNA molecules were performed in a multichannel microfluidic system (Lumicks). Bead positions and applied tension were controlled with a 30 ms resolution using custom LabView software.

Fluorescent images of dCasCas12d-sgRNA-DNA complexes were collected with an EMCCD camera (Cascade II, Photometrics) using the Micromanager software. Fluorescence light was excitated with a DPSS 473 nm CW laser (100 mW, Lasever, LSR473U) and separated using filter set 10 (Carl Zeiss). Images were obtained with an exposure time of 1000 ms and electron multiplier gain of 3800. Saved TIFF files were further processed using FIJI ImageJ.

### Single-molecule assay

Double-stranded DNA molecules (48.5 kbp) with biotinylated ends were prepared from bacteriophage lambda DNA, as described previously [15]. For single-molecule experiments 100 nM dCas12d E913A was individually mixed with 200 nM of RNA1/2/3/4 in four separate tubes in 1x CutSmart buffer at room temperature. After a short incubation at room temperature the four solutions were mixed together. All experiments were performed in 40 mM Tris-OAc (pH = 8.0), 80 mM KOAc, 10 mM MgOAc, and 0.02% BSA at 37°C. The inner surfaces the of microfluidic system were passivated with 0.5% Pluronic F-127 and 0.1% BSA [17]. Two channels of microfluidic cell were used to obtain single optically trapped DNA tether and thus were fed with a 0.01% solution of a 2.1 μm streptavidin-coated polystyrene beads (Spherotech) and 1.25 pM solution of a double stranded DNA with biotinylated ends. After the DNA was captured, its force-extension behavior was measured to verify that a single DNA molecule was tethered between the two beads. Next, the DNA tether was incubated in a channel containing 100 nM dCas12d-sgRNA for 3 min and then was fluorescently visualized in a protein-free channel.

## RESULTS

### Production of catalytically inactive versions of the KbCas12d effector

Bioinformatic analysis of the CRISPR-KdCas12d locus from shows that the system includes a 1041 bp *cas1* gene encoding protein and a *kbCas12d* effector gene of 3378 bp.[7] The system includes 14 spacers of 17-18 nucleotides in length separated by repeat sequences of 26 nucleotides in length. A sequence encoding scoutRNA is located upstream of the CRISPR array. Transcription of the scoutRNA and CRISPR-RNA is in the same direction (Figure 1A).[7,11]

**Fig. 1.**
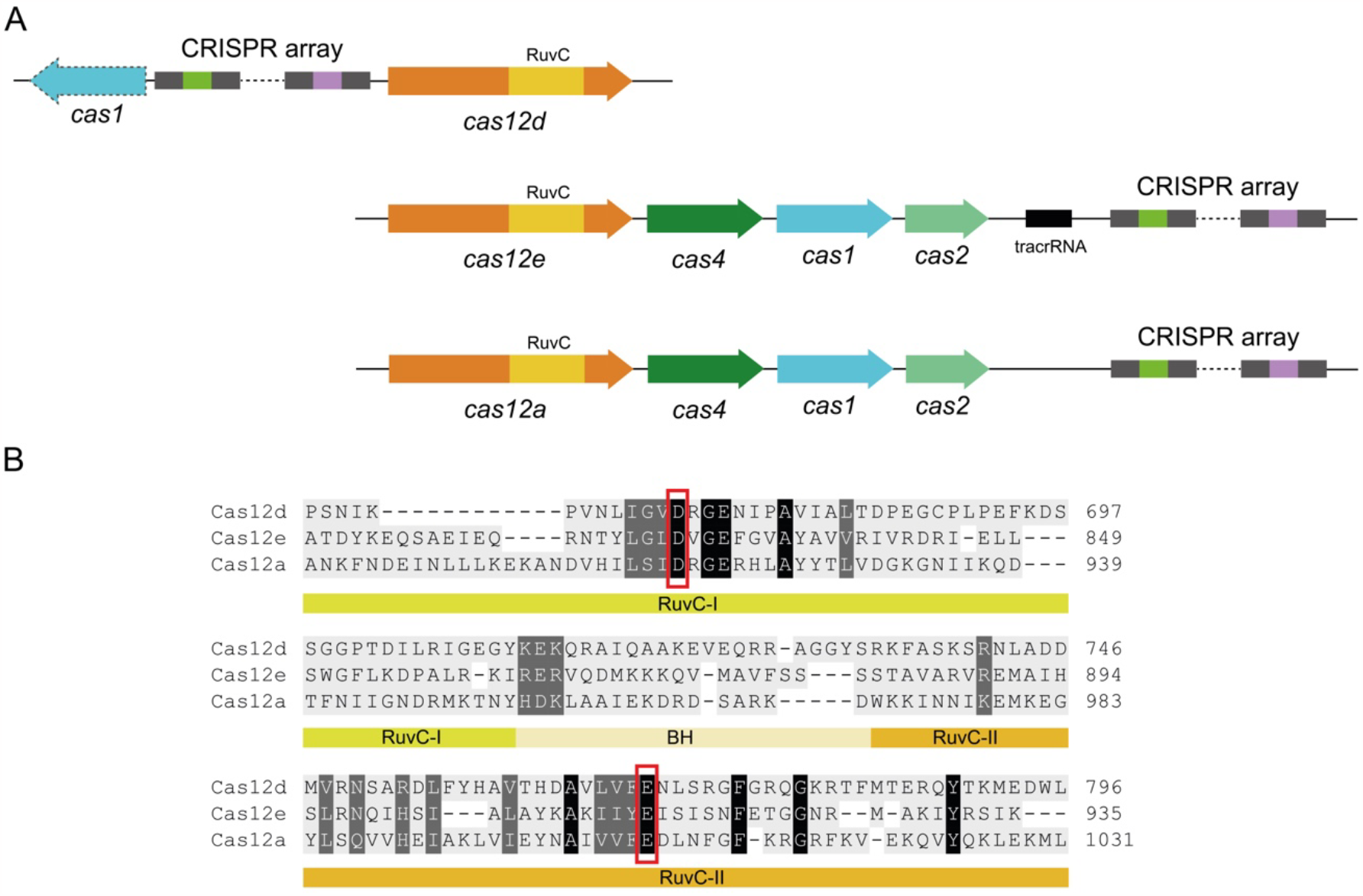
(**A**) Organization of type V CRISPR-Cas systems: Cas12d, Cas12a and Cas12e. (**B**) Fragment of the KbCas12d, DpbCas12e and AsCas12a RuvC domains alignment. Known mutation sites for the AsCas12a protein are D917A and E1006A. Known mutation sites of DpbCas12e protein: D672A and E769A. Corresponding KbCas12d protein mutation sites: D827A and E913A.

To obtain catalytically inactive forms of the effector KbCas12d, its amino acid sequence was analyzed to identify possible functional domains. Using catalytically inactive mutant data for CRISPR-Cas12e [12] and CRISPR-Cas12a [13], the identified RuvC nuclease domains of all three systems were aligned. Resulting alignment (Figure 1B) suggested substitutions in the amino acid sequence of the RuvC domain of the KbCas12d system at positions D827 and E913 for alanine to create “dead” forms – dKbCas12d (DNA-binding but non-cutting versions of the protein).

The cloning and purification of mutant and wild-type proteins are described in the Methods section. As a result, recombinant proteins were successfully obtained in monomeric form at sufficient concentrations.

### Design of single guide RNA

In order to improve the cleavage activity, an optimal scoutRNA was designed. For this purpose, a number of scoutRNA (1-5) forms differing in length and secondary structure were synthesized based on scoutRNA sequence data [11](sequences are given in Supplementary information). *In vitro* reactions contained 5 types of different synthesized scoutRNA, crRNA, recombinant KbCas12d protein and target DNA (818 b. p.) were performed separately in a 20 μl volumes at 37°C for 30 minutes (1X CutSmart buffer (NEB), 20 nM DNA, 4 μM scoutRNA/crRNA, 400 nM KbCas12d protein) The experimental results (Figure 2A) indicated that scoutRNA1, the shortest form of the tested molecules, was optimal for DNA cleavage. This scoutRNA1 was used as the basis for the design of a hybrid RNA, a fusion molecule of guide crRNA and scoutRNA (sgRNA, single guide RNA, sgRNA) (Figure 2C). To create sgRNA, the 3’-end of the scoutRNA1 was joined to the crRNA sequence via linkers by the “GAAA” sequence. sgRNAs with 3 different linker lengths were tested (Figure 2B and 2C): 400 nM KbCas12d protein in complex with 4 μM sgRNA1, sgRNA2 or sgRNA3 was incubated with the DNA target used in the above experiments for 30 min at 37°C. The sample contained a mix of crRNA and scout RNA was used as the positive control. The reaction products were applied to agarose gel electrophoresis to evaluate the cleavage efficiency. The results of the experiment demonstrated (Figure 2B) that the use of sgRNA1, sgRNA2 led to even more efficient cleavage of target DNA, compared to mix scout and crRNA. sgRNA2 has been chosen for further research. Thus, it was possible to identify a form of hybrid RNA (sgRNA) for efficient DNA cleavage by the KbCas12d protein (Figure 2C). The algorithm used for scoutRNA and crRNA joining could potentially be used for other representatives of the Cas12d family.

**Fig. 2.**
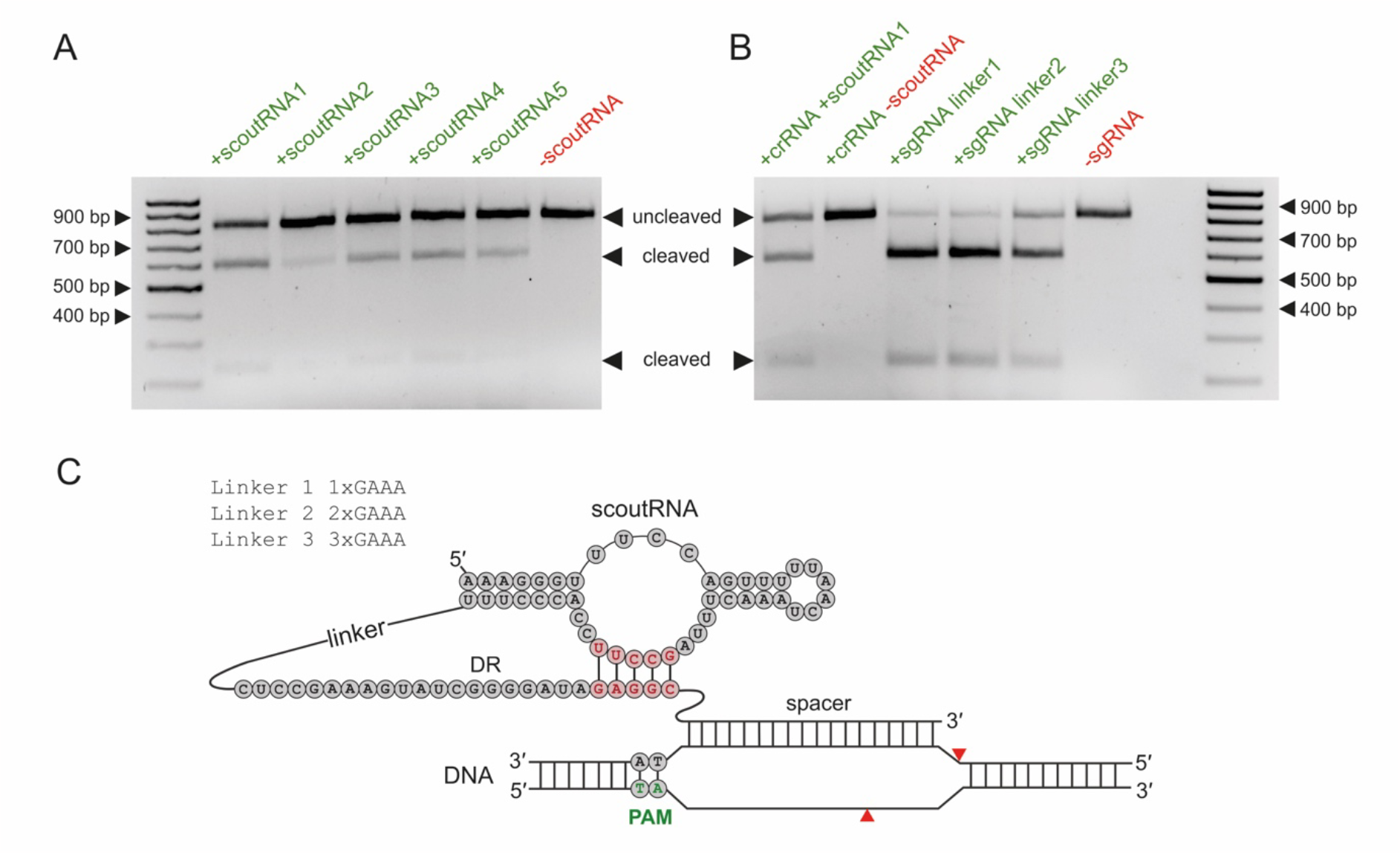
(**A**) *In vitro* testing of scoutRNAs with different lengths (**B**) *In vitro* testing of sgRNAs with different linker lengths. Reaction containing a mix of crRNA and scoutRNA was used as a positive control. Black arrows indicated the cleavage fragments. (**C**) Design of the KbCas12d sgRNA binding to target DNA

### The nuclease and binding activity of dKbCas12d D827A and dKbCas12d E913A

The absence of nuclease activity of the mutant forms of KbCas12d was tested. *In vitro* DNA cleavage reaction was performed with ratios of target DNA to ribonucleic acid complex (RNP) (1:1, 1:10, and 1:20); samples were incubated under the conditions described above for 30 min at 37°C. The reaction was then split into equal parts, and the samples were loaded on a native agarose gel (Figure 3A) and 5% PAGE under denaturing conditions (Figure 3B) to demonstrate the absence of DNA nicking (Figure 3B). As a result, it was observed that wild-type KbCas12d performs targeted double-stranded break whereas mutants are incapable of any nuclease activity.

**Fig. 3.**
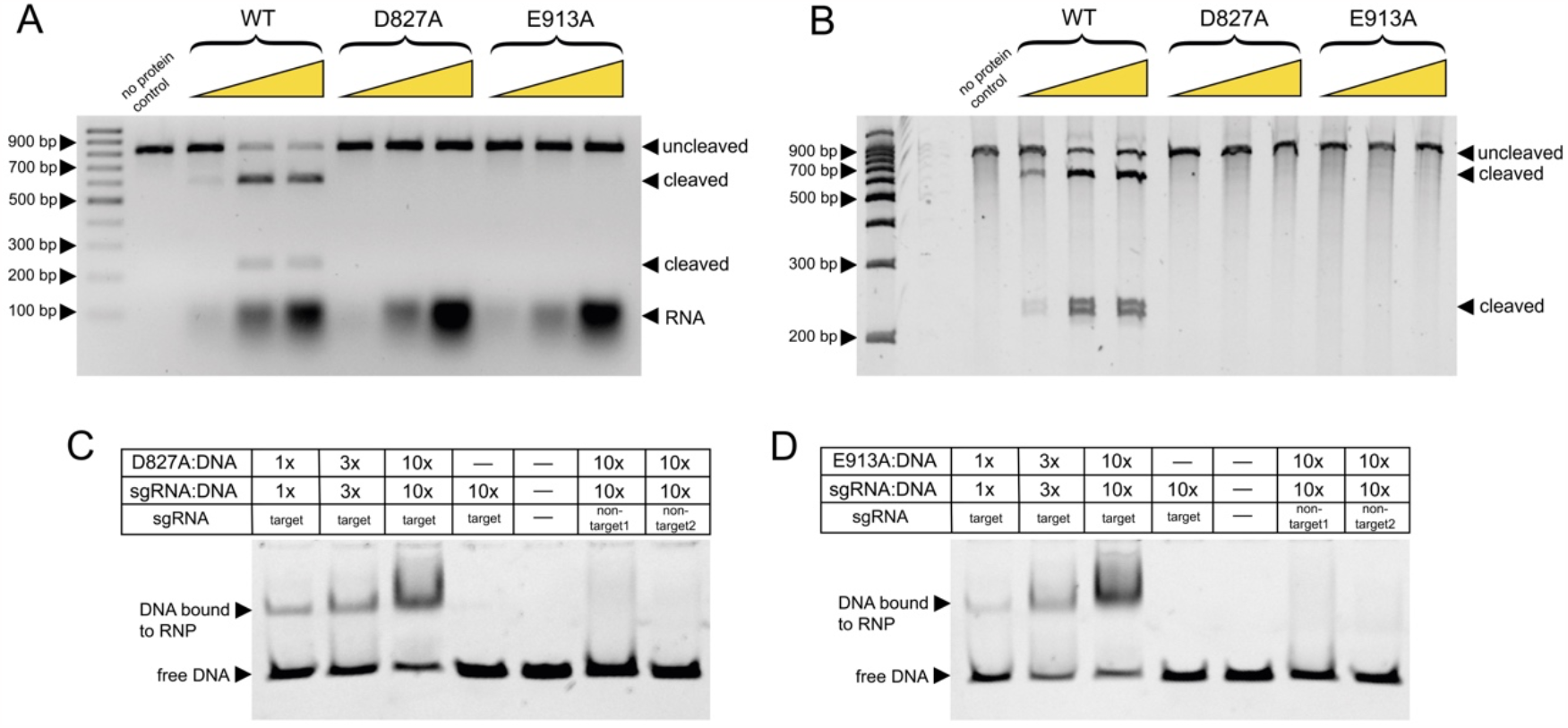
(**A**) Nuclease activity of wild-type KbCas12d and mutant dKbCas12d D827A and dKbCas12d E913A RNP complexes incubated at different ratios with target DNA (1:3, 1:10, and 1:20). Agarose gel electrophoresis under native conditions. The fragments produced as a result of the cleavage are marked with black arrows (**B**) Nicking activity of wild-type KbCas12d and mutant dKbCas12d D827A and dKbCas12d E913A RNP complexes incubated at different ratios with target DNA (1:3, 1:10, and 1:20). Polyacrylamide gel electrophoresis under denaturing conditions. (**C, D**) Binding of an RNP complexes of the dKbCas12d D827A and dKbCas12d E913A to the target DNA.

Next, the ability of the dKbCas12d D827A and dKbCas12d E913A mutants to bind to the target DNA was tested (Figure 3C). KbCas12d mutants in complex with sgRNAs was mixed with the Cy-3 tagged DNA target *in vitro* at protein/RNA:DNA ratios of 1:1, 1:3, and 1:10 (see Methods section). After 30 min incubation at 37°C the samples were loaded onto a 5% polyacrylamide gel and electrophoresis was run on an ice bath. The EMSA assay results (Figure 3C) demonstrated changes in the mobility of labeled Cy-3 DNA in complex with target sgRNA and mutant KbCas12d proteins. In samples where non-target RNAs were used, there was no shift observed. Thus, the obtained variants of the dKbCas12d D827A and dKbCas12d E913A are able to bind specifically to the target DNA and cannot perform a cleavage.

### Specificity of catalytic inactive KbCas12d binding to DNA at the single molecule level

To assess the efficiency and specificity of DNA binding to extended molecules, we used a single-molecule laser tweezers approach. For this experiment, we chose the dKbCas12d E913A version. We designed four sgRNAs containing FAM-tagged UTPs targeting four equally spaced (approximately 5000 bp) regions of lambda phage DNA (RNAs listed in Supplementary file). Then, we used a custom optical trapping system and a microfluidic chamber, (see Methods section) to capture lambda DNA between two microspheres and incubated for 3 min in a channel containing a mixture of four RNP, keeping the DNA between the traps in a relaxed state (Figure 4 A). We then transferred the system to a channel containing only the buffer and performed fluorescence imaging (Figure 4B). We observed (n=20 DNA molecules) that RNP complexes were often bound to specific target sites, albeit with low varying efficiency. Only in a few of cases we were able to observe simultaneous landing of three RNP complexes at the same time (Figure 4 B, C). Since the amount of fluorescent UTPs can vary between molecules, we observed different signal intensities from the targets (Figure 4C).. Data analysis revealed nearly equal binding probability of RNP to four specific targets T1/T2/T3/T4: 0.19 ± 0.1, 0.25 ± 0.18, 0.33 ± 0.17 and 0.28 ± 0.17 (mean ± SEM). Almost no non-specific interactions were observed during the experiments.

**Fig. 4.**
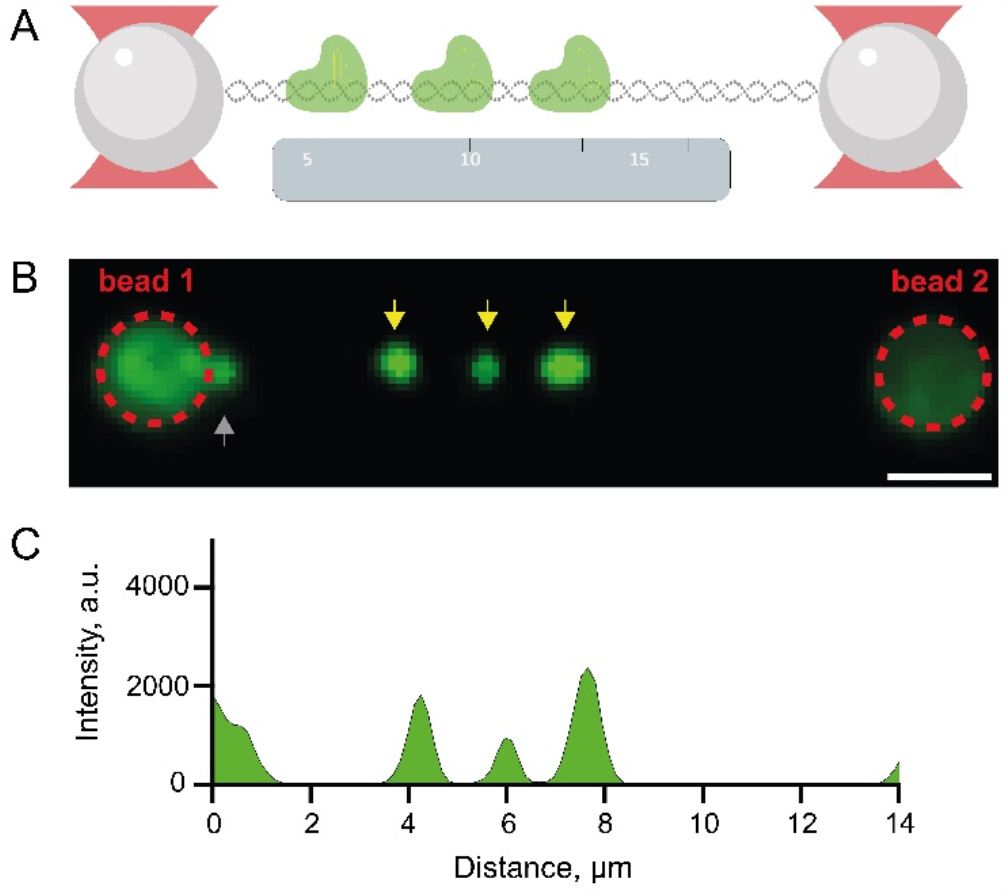
Single-molecule visualization of dKbCas12d E913A -sgRNA-DNA complexes. (**A**) Schematic of experimental workflow. (**B**) Representative fluorescent image, revealing three specific complexes (yellow arrows) and non-specific binding (gray arrow) on 48.5 kbp DNA containing four targets. (**C**) Corresponding intensity plot profile.

## DISCUSSION

This study performs new catalytically inactive mutants of the Cas12d system, which are unable to make DNA double-stranded breaks, but have the ability to bind to the target DNA. Despite the significant differences in organization, the structure of the Cas12d RuvC nuclease domain aligned well with the amino acid sequences of the Cas12e and Cas12a [5,12,13,18,19]effectors. The structure of the protein itself may be closer to that of Cas12e, however this requires further structural studies. On the other hand there is a significant difference in the CRISPR-RNA structure, where a characteristic scoutRNA should be present for the CRISPR-Cas V-D system. Although the rather long and conserved scoutRNAs were described in the literature [11]we have shown that significantly shortened versions also exhibit *in vitro* activity. Therefore, the identification of all features of the scout RNA remains to be elucidated. Nevertheless, we were able to create an efficient single-guide RNA in which both of crRNA and scoutRNA are bound through a linker and demonstrate its ability to accomplish directed double-strand breaks for a wild-type system. Such a catalytically active effector-based system could be used for genetic editing in the future.

Using standard biochemical methods, we demonstrated the ability of the dKbCas12d D827A and dKbCas12d E913A mutants to bind target DNA using a single guide RNA. Subsequently, we employed optical tweezers to evaluate the specificity and efficiency of the dKbCas12d E913A system at the single-molecule level. Our data showed good specificity, which is similar to other single-molecule research on Cas12a [20]. Also, according to published data [21], Cas12a is more sensitive than Cas9 to the introduction of artificial mismatches in the guide RNA (16 vs. 9 PAM proximal matches). The Cas12d system is characterized by a spacer sequence comprised of 17-18 nucleotides. Substituting any of these bases may significantly impact the recognition and nuclease activity of the complex [11]. Thus, the low binding efficiency of the complexes and the lack of off-target binding may be explained by the fact that a prolonged binding event, which we are able to capture, requires the complementarity of all spacer nucleotides. Low off-target binding efficiency appears because of the complex’s rapid and specific DNA molecule scanning.

Many mechanisms of the CRISPR KbCas12d system remain to be thoroughly elucidated, but its unique features indicate its high potential for future use in diagnostic, medical and biotechnological applications.

